# Differential TAM receptor regulation of hepatic physiology and injury

**DOI:** 10.1101/2020.03.13.990143

**Authors:** Anna Zagórska, Paqui G. Través, Lidia Jiménez-García, Jenna D. Strickland, Francisco J. Tapia, Rafael Mayoral, Patrick Burrola, Bryan L. Copple, Greg Lemke

**Author notes:** These authors contributed equally to this work (Anna Zagórska, Paqui G. Través). AZ and PGT are currently employees of Gilead Sciences, Inc., and RM is currently an employee of Merck & Co. All authors declare no conflicts of interest. Correspondence: Greg Lemke, The Salk Institute for Biological Studies, 10010 North Torrey Pines Road, La Jolla, CA, 92037.

## Abstract

The TAM receptor tyrosine kinases (RTK) Mer and Axl have been implicated in liver disease, yet our understanding of their roles in liver homeostasis and injury is limited. We therefore examined the performance of Mer and Axl mutant mice during aging, and in four models of liver injury. We find that Mer and Axl are most prominently expressed in Kupffer and hepatic endothelial cells, and that as *Axl*^*-/-*^*Mertk*^*-/-*^ mice normally age, they develop profound liver disease. We further find that Mer signaling is critical to the phagocytosis of apoptotic hepatocytes that are generated during acute hepatic injury, and that Mer and Axl act in concert to inhibit injury-triggered cytokine production. TAM expression in Kupffer cells is crucial for these effects. In contrast, we show that Axl is uniquely important in mitigating liver damage during acute acetaminophen intoxication. Finally, we demonstrate that Axl exacerbates the fibrosis that develops in a model of chronic hepatic injury. These divergent effects have important implications for the design and implementation of TAM-directed therapeutics that target these RTKs in the liver.

## Introduction

Liver diseases - including acute liver failure, viral hepatitis, and alcoholic and non-alcoholic fatty liver disease (NAFLD) – represent a major medical burden worldwide (1-3). Increasing evidence suggests that both progression and resolution of these diseases depends on the kinetics and intensity of innate and adaptive immune responses (4, 5); and that macrophages - including Kupffer cells (KCs), the resident macrophages of the liver – are important regulation loci (6).

We have shown that the TAM RTKs - Tyro3, Axl, and Mer (7) – are pivotal modulators of tissue macrophage function generally (8-13). Over the last several years, genome-wide association studies have tied polymorphisms in the human *MERTK* gene - encoding Mer - to altered risk for both (a) fibrosis in patients with chronic Hepatitis C virus (HCV) infection (14-17), and (b) NAFLD, in which two intronic single-nucleotide *MERTK* polymorphisms are protective (18, 19). In the progression from NAFLD to nonalcoholic steatohepatitis (NASH), these polymorphisms, which are associated with *reduced* Mer expression, are linked to reduced risk for liver fibrosis (20). In turn, recent analyses have indicated that *Mertk*^*-/-*^ mice display reduced levels of a NASH-like fibrosis that is induced by high-fat diet, via reduced activation of hepatic stellate cells by macrophages that are normally Mer^+^ (21). Together, these findings suggest that Mer signaling promotes hepatic fibrosis. Independently, patients with acute liver failure have been found to display markedly elevated numbers of Mer^+^ macrophages and monocytes in their liver, lymph nodes, and circulation (22-24), and Mer has therefore emerged as a target in the treatment of liver disease (25, 26).

With respect to Axl, elevated serum levels of soluble Axl extracellular domain (sAxl) have been found to be a biomarker for hepatocellular carcinoma (27), and mice lacking Gas6, the obligate Axl ligand (28), display enhanced tissue damage in a liver ischemia model (29). At the same time, Axl^+^ monocytes are elevated in patients with cirrhosis (30), and serum Gas6 and sAxl levels are elevated in patients with HCV and alcoholic liver disease (24). Divergent roles for Axl and Mer have been reported in chronic models of fibrosis, where *Mertk*^*-/-*^ mice exhibited enhanced NASH development when fed a high fat diet, while *Axl*^*-/-*^ mice were protected (31). These multiple findings notwithstanding, the general importance of TAM receptor signaling to both normal liver physiology and to acute, rapid-onset liver insults has not been assessed. We have therefore exploited a set of conventional and conditional mouse mutants in the *Axl* and *Mertk* genes, and subjected these mutants to established models of both acute liver damage and chronic fibrosis, in order to make these assessments.

## Materials and Methods

### Mice

The *Axl*^−/−^, *Mertk*^−/−^, *Axl*^−/−^*Mertk*^*-*–/–^, TAM TKO, *Cx3cr1*^*GFP/+*^*Axl*^*-/-*^*Mertk*^*-/-*^ and *Cx3cr1*^*Cre/+*^*Axl*^*f/f*^*Mertk*^*f/f*^, *Cx3cr1*^*CreER/+*^*Axl*^*f/f*^*Mertk*^*f/f*^ mice were as described previously (9, 32-34). All lines, except for the TAM TKOs, have been backcrossed for >9 generations to a C57BL/6 background. All animal procedures were conducted according to guidelines established by the Salk Institutional Animal Care and Use Committee. Mice were randomly allocated to experimental groups. Mice were fed irradiated rodent diet 15053 (Lab Diet), caged in individual ventilated cages with Anderson 0.25 inch corn cob bedding, and maintained on a 12 hour light-dark cycle.

### Acute liver damage models

In the fulminant hepatitis (Jo2) model, mice were injected with an anti-CD95 monoclonal antibody (Jo2 clone; BD Pharmingen Cat# 554254), which activates the Fas receptor and induces apoptosis of hepatocytes (35, 36). For a lethal dose, 8-12 week old mice (males and females) were injected IP with 1 mg/kg body weight of anti-mouse CD95 antibody (Jo2) and monitored every 30 min. For a sub-lethal dose, 8-12 week old mice were injected IP with 0.3 mg/kg Jo2 antibody. The D-galactosamine (D-gal)/lipopolysaccharide (LPS) model mimics bacterial peritonitis and endotoxic shock (37). For a sub-lethal dose, 8-12 week old mice (males and females) were injected IP with D-gal (Sigma Aldrich, Cat# G0500, 350 mg/kg body weight) and LPS (ENZO life sciences, Cat# ALX-581-013-L002, 1 μg/kg body weight) in saline (final volume 0.3-0.4 ml). Mice were sacrificed at the indicated times post injection. Serum and liver samples were collected for histological and biochemical analysis.

For the acetaminophen toxicity model, mice were treated with 300 mg/kg acetaminophen (10 μl/g) (Sigma Chemical Company) in sterile saline, or sterile saline vehicle alone, by intraperitoneal injection. Mice were fasted in clean cages for 16 hours (overnight) prior to injection the following morning. (Fasting is necessary to produce acetaminophen-induced liver injury in mice.) Mice were monitored for general appearance, motility, and survival at 12, 24, and 48 hours after treatment. While the goal of these studies was to evaluate the impact of TAM deletion on the resolution of inflammation, which occurs between 48 and 72 hours after acetaminophen overdose (38), this was precluded by the severe phenotypes that appeared in the *Axl*^*-/-*^ mice as soon as 12 hours after treatment. Liver and blood were collected from mice at 48 hours after acetaminophen. At 48 hours, mice were anesthetized with pentobarbital (50 mg/kg) and euthanized by exsanguination.

### Carbon tetrachloride liver fibrosis model

Carbon tetrachloride (CCl4; Sigma) was injected IP at 0.5 ml/kg, diluted in corn oil (Sigma) at 0.2 ml final volume, three times per week for 6 weeks. (39). Two days after the last injection, mice were euthanized and serum and liver were collected for histological and biochemical analysis.

### Antibodies

Antibodies used were: Mer for WB (R&D AF591), Mer for IHC (eBioscience DS5MMER), Axl for IHC (R&D, AF854), Axl for immunoprecipitation (Santa Cruz, M-20), GAPDH (Millipore, MAB374, clone 6C5), phospho-tyrosine (Millipore, clone 4G10), Gas6 (R&D, AF986), CD31 (Abcam, ab28364), MARCO (AbD Serotec MCA1849), Laminin (Sigma L9393), Desmin (Abcam ab32362), cleaved Caspase 3 (Asp175, Cell Signaling), F4/80 (AbD Serotec MCA497), Ki67 (Biolegend 65241) GFAP (Dako, z0334). Axl, Mer, and Gas6 antibody specificity for immunohistochemistry, immunocytochemistry, immunoprecipitation and immunoblotting was tested using samples from corresponding mouse mutants (for example, Fig. 1G, H). Secondary antibodies used for immunoblot analysis were horseradish-peroxidase-conjugated anti-goat (705-035-003) from Jackson ImmunoResearch, and anti-mouse (NA931V) and anti-rabbit (NA934V) from GE Healthcare. Secondary antibodies for immunocyto and immunohistochemistry were fluorophore-conjugated anti-goat (A-11055 from Life Technologies, or 705-166-147 from Jackson ImmunoResearch), anti-rabbit (A-10040 or A-21206 from Life Technologies), anti-rat (712-545-153 or 712-165-153 from Jackson ImmunoResearch), and anti-mouse (A-11029 from Life Technologies, or 715-166-150 from Jackson ImmunoResearch).

**FIG. 1.**
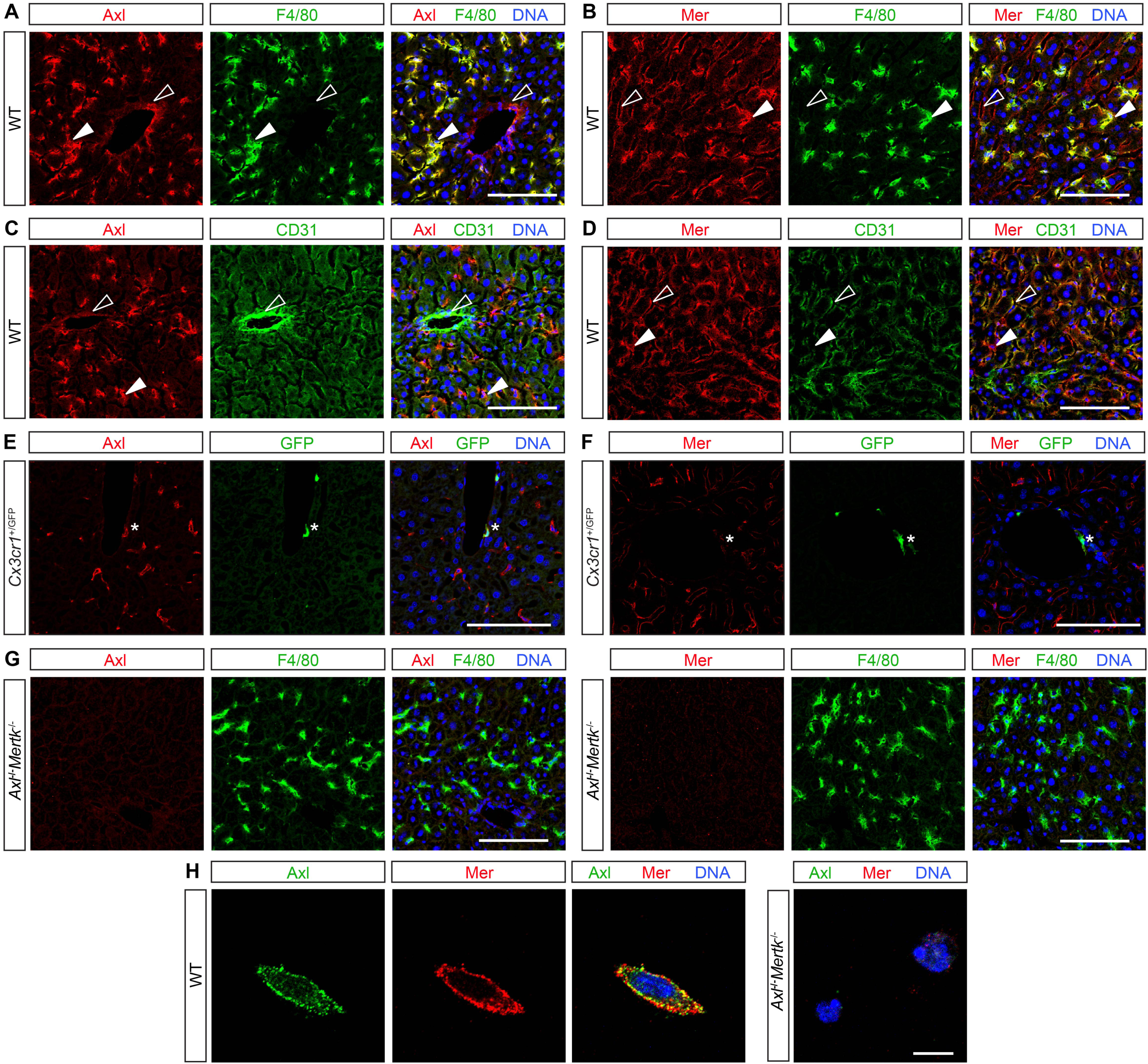
TAM expression in normal liver. (A-G) Mouse liver immunohistochemistry. (A,C,E) Axl (red) is expressed by F4/80^+^ KCs (green, closed arrowheads) and strongly CD31^+^ blood-vessel-lining endothelial cells (green, open arrowheads) and by perivascular macrophages (E, green, asterisk). (B,D,F) Mer (red) is expressed by KCs (green, closed arrowheads), by weakly CD31^+^ sinus-lining endothelial cells (green, open arrowheads), and very weakly by perivascular macrophages (F, green, asterisk). (G) *Mertk*^-/-^*Axl*^-/-^ mice were used for antibody control. (H) Basal TAM expression on isolated KCs. Axl and Mer are co-expressed on the surface of individual cultured KCs. (See methods for KC purification and culture.) Bars A-G, 100 µm; H, 10 µm.

### Kupffer cell isolation

Non-parenchymal cells (NPCs) were prepared by two-step *in situ*/*ex situ* collagenase/pronase digestion and fractionation on a continuous density gradient of 36% Percoll prepared in GBSS/B (Gey’s Balanced Salt Solution supplemented with 0.8% NaCl at pH 7.35). The procedure was carried out in anesthetized animals (100:10 mg/kg Ketamine:Xylazine). Perfusion was done *in situ* via portal vein with a 24Gx3/4” catheter. Hanks balanced salt solution (HBSS) without Ca^2+^ was perfused at 37°C with a flow rate of 10 ml/min. During the first 5 min of perfusion, regular HBSS was used as a washing solution. Subsequently, HBSS containing 0.04% Collagenase Type 2 (Worthington), 50 mM HEPES (N-2-hydroxyethylpiperazine-N’-2-ethanesulfonic acid), and 0.6 mg/ml CaCl_2_*2H_2_O was substituted and the perfusion continued during 5 min. The perfused liver was excised, minced *ex situ* and mixed with 30 ml of HBSS containing 0.07% Pronase E (Merck EMD), 50 mM HEPES and 0.6 mg/ml CaCl_2_*2H_2_O. 20 min after continuous stirring at 37°C, the cell suspension generated was filtered through a 100µm cell strainer (Falcon), and the filtrate was centrifuged three times for 5 min (50 x g) in 50 ml of HBSS supplemented with 0.2mg/ml EGTA and 20µg/ml DNAse-I, to pellet the hepatocytes. The three supernatants obtained were used for the preparation of Kupffer cells by centrifugation over a 36% Percoll continuous density gradient (density, 1.066 g/ml; pH 7.5 in GBSS/B). Hepatocyte-depleted supernatants were subjected to a second centrifugation (600 x g, 4°C, 5 min), and the pellets were mixed, resuspended with 40 ml Percoll-GBSS/B Solution (36% Percoll), transferred in one 50 ml tube, adding 500µl DNAse-I (2mg/ml) and centrifuged in a swinging bucket rotor during 20 min (800 x g, 4°C) without brakes. To remove the Percoll, the pellet was resuspended in 14 ml of GBSS/B and centrifuged 5 min (800 x g, 4°C). The pellet was resuspended in 4 ml of Red Blood Lysis Buffer during 10 min (eBioscience 00-4333-5710) to remove erythrocyte contamination and centrifuged 5 min (800 x g, 4°C). The final precipitate is an enriched fraction containing NPCs, but it is practically free of debris and hepatocytes. KCs were selectively removed from the fractions by selective attachment to plastic. The resuspended NPC pellet was plated at 10×6 cells per cm^2^ in six-well cluster dishes (Becton Dickinson) and cultured in RPMI 1640 supplemented with 10% fetal calf serum (FCS) and antibiotics (50 µg/ml each of penicillin, streptomycin and gentamicin). KCs were allowed to attach to the plastic for 30 min and were washed several times with PBS to remove unattached cells.

### Immunoblotting and immunoprecipitation

Thioglycolate-elicited peritoneal macrophages were washed with ice-cold DPBS and lysed on ice in a buffer containing 50 mM Tris-HCl pH 7.5, 1 mM EGTA, 1 mM EDTA, 1% Triton X-100, 0.27 M sucrose, 0.1% β-mercaptoethanol, and protease and phosphatase inhibitors (Roche). Tissues were snap frozen in liquid nitrogen prior to lysis. For immunoblots, equal amounts of protein (10 μg) in LDS sample buffer (Invitrogen) were subjected to electrophoresis on 4–12% Bis-Tris polyacrylamide gels (Novex, Life Technologies) and transferred to PVDF membranes (Millipore). For immunopreciptations, cell lysates were incubated overnight (ON) at 4°C with indicated antibodies. Protein G-Sepharose (Invitrogen) was added for 2 h and immunoprecipitates (IPs) were washed twice with 1 ml of lysis buffer containing 0.5 M NaCl and once with 1 ml of 50 mM Tris-HCl pH 7.5. IPs were eluted in LDS buffer, separated on polyacrylamide gels and transferred to PVDF membranes. Membranes were blocked in TBST (50 mM Tris-HCl pH 7.5, 0.15 M NaCl, and 0.25% Tween-20) containing 5% BSA and immunoblotted ON at 4°C with primary antibodies diluted 1000-fold in blocking buffer. Blots were then washed in TBST and incubated for 1 h at 22-24°C with secondary HRP-conjugated antibodies (GE Healthcare) diluted 5000-fold in 5% skim milk in TBST. After repeating the washes, signal was detected with enhanced chemiluminescence reagent.

### sAxl, TNF alpha, and IL-1 beta ELISA assays

ELISA for measurement of sAxl, Gas6 (R&D), TNF alpha, and IL-1 beta (eBiosciences) were performed according to manufacturers’ instructions.

### ALT and AST assay

ALT (TR71121) and AST (TR70121) assays were from Thermo Scientific, and were performed according to manufacturer instruction.

### Sirius red stain

Sections were fixed for 24h in PFA 4% at RT, washed in PBS two times 5 min and stained with Hematoxylin for 8 min at RT. Slides were then washed twice in warm water for 5 min and stained for 1h at RT with Picro-Sirius Red solution (0.1% (m/v) Sirius Red (Sigma) in saturated aqueous solution of Picric Acid (LabChem)). Slides were washed two times 5 min with “acidified water” (acetic acid 0.5%) and dehydrated in 100% EtOH three times 2 min. Finally, the slides were cleared in Histo-Clear (National Diagnostics) for 5 min and mounted with VectaMount (Vector Labs).

### Immunocytochemistry and immunohistochemistry

For immunohistochemistry, tissues were fresh frozen and cut into 11 μm sections, air-dried and stored desiccated at −70°C. Prior to staining, sections were fixed for 3 min with ice-cold acetone, washed in PBS, blocked in blocking buffer (PBS containing 0.1% Tween-20, 5% donkey serum and 2% IgG-free BSA) for 1 h. Slides were then washed in PBS 0.1% Tween-20 and incubated with 1 μg/ml primary antibody in blocking buffer ON at 4°C. Slides were washed five times 5 min in PBS 0.1% Tween-20 and incubated with Hoechst and fluorophore-coupled donkey (Jackson) secondary antibodies diluted 1:400 in blocking buffer for 2 h at 22-24°C in dark. Slides were washed, sealed with Fluoromount-G (SouthernBiotech) and stored at 4°C. For immunocytochemistry, cells were first fixed in 4% PFA for 10 min and washed with PBS. Cover slips were then blocked for 30 min in blocking buffer with 0.1% Triton X-100, washed in PBS 0.1% Tween-20 and incubated with primary antibody for 1 h at 22-24°C. Cover slips were washed five times in PBS 0.1% Tween-20 and incubated with Hoechst and fluorophore-coupled donkey secondary antibody diluted 1:400 in blocking buffer for 1 h at 22-24°C in dark. Cover slips were washed and mounted on slides with Fluoromount-G. Images were taken on a Zeiss LSM 710 microscope with Plan-Apochromat 20x/ 0.8 M27 and 63x/ 1.40 Oil DIC M27 objectives. Morphometric quantification was performed using Image J to quantify % marker area or number of positive cells per area. 10-15 images from 3 separate sections per mouse were quantified.

### RT-qPCR

Total cellular RNA was isolated using TRIzol Reagent (Thermo), according to manufacturer’s instructions. Reverse transcription was performed with RT Transcriptor First Strand cDNA Synthesis Kit (Roche) with anchored oligo-dT primers (Roche). qPCR was run in a 384-well plate format on a ViiA 7 Real-Time PCR System (Applied Biosystems) using 2x SYBR Green PCR Master Mix (Applied Biosystems). Analysis was done using delta delta Ct method. Primers are listed in **Supporting Table 1**. Cyclophilin A (Ppia) and 36B4 (Rplp0) were used as control housekeeping genes.

### Data analysis

Statistical analysis was performed using two-tailed Student’s *t*-test.

## Results

### Expression of TAM receptors in mouse liver

We first used immunohistochemistry (IHC) to delineate TAM expression in adult mouse liver. Most prominently, we detected very strong expression of both Axl (Fig. 1A) and Mer (Fig. 1B) in all KCs. These liver macrophages did not express detectable Tyro3 (data not shown). Most tissue macrophages express high and low levels of Mer and Axl, respectively, and so KCs fall into the restricted subset of macrophages, including red pulp macrophages of the spleen (40), that abundantly express both. Axl and Mer were also expressed in many CD31^+^ endothelial cells of the liver vasculature, with Axl in strongly CD31^+^ blood vessels (Fig. 1C), and Mer in weakly CD31^+^ hepatic sinuses (Fig. 1D). Axl was also expressed in perivascular macrophages, which only weakly expressed Mer (Fig. 1E,F). Antibody specificity was confirmed using *Axl*^*-/-*^*Mertk*^*-/-*^ tissues and cells (Fig. 1G,H). Axl and Mer were co-expressed on freshly isolated KCs (Fig. 1H).

### Role of TAM receptors in hepatic aging

Given these expression data, we asked if Axl and Mer were relevant to normal hepatic aging by examining the livers of aged (7-12 month) *Axl*^*-/-*^*Mertk*^*-/-*^ mice versus wild-type (WT).

Remarkably, we found that multiple inflammation and tissue damage markers were markedly elevated in the aged *Axl*^*-/-*^*Mertk*^*-/-*^ liver, in the absence of any overt perturbation (Fig. 2). Cleaved Caspase 3^+^ (cCasp3^+^) apoptotic cells (ACs) were elevated 10-fold (Fig. 2A), consistent with the essential role that TAM receptors play in AC clearance (10, 13, 28, 41). Liver expression of the scavenger receptor MARCO, an indicator of macrophage activation, was dramatically higher in the double mutants (Fig. 2B); as was GFP expression in *Cx3cr1*^*GFP/+*^*Axl*^*-/-*^*Mertk*^*-/-*^ mice, indicative of the immune infiltration of CX3CR1^+^ monocytes (Fig. 2C). Liver mRNAs encoding multiple proinflammatory cytokines, immune modulators, and chemokines were elevated 4-to 20-fold in aged *Axl*^*-/-*^*Mertk*^*-/-*^ mice compared to WT (Fig. 2D), and the serum levels of two enzymes associated with liver damage – alanine and aspartate aminotransferase (ALT and AST, respectively) – were similarly elevated (Fig. 2E). Together, these results demonstrated that the aged *Axl*^*-/-*^*Mertk*^*-/-*^ livers were damaged and inflamed in the absence of any experimental insult, and that TAM signaling is therefore required for normal liver homeostasis and healthy aging.

**FIG. 2.**
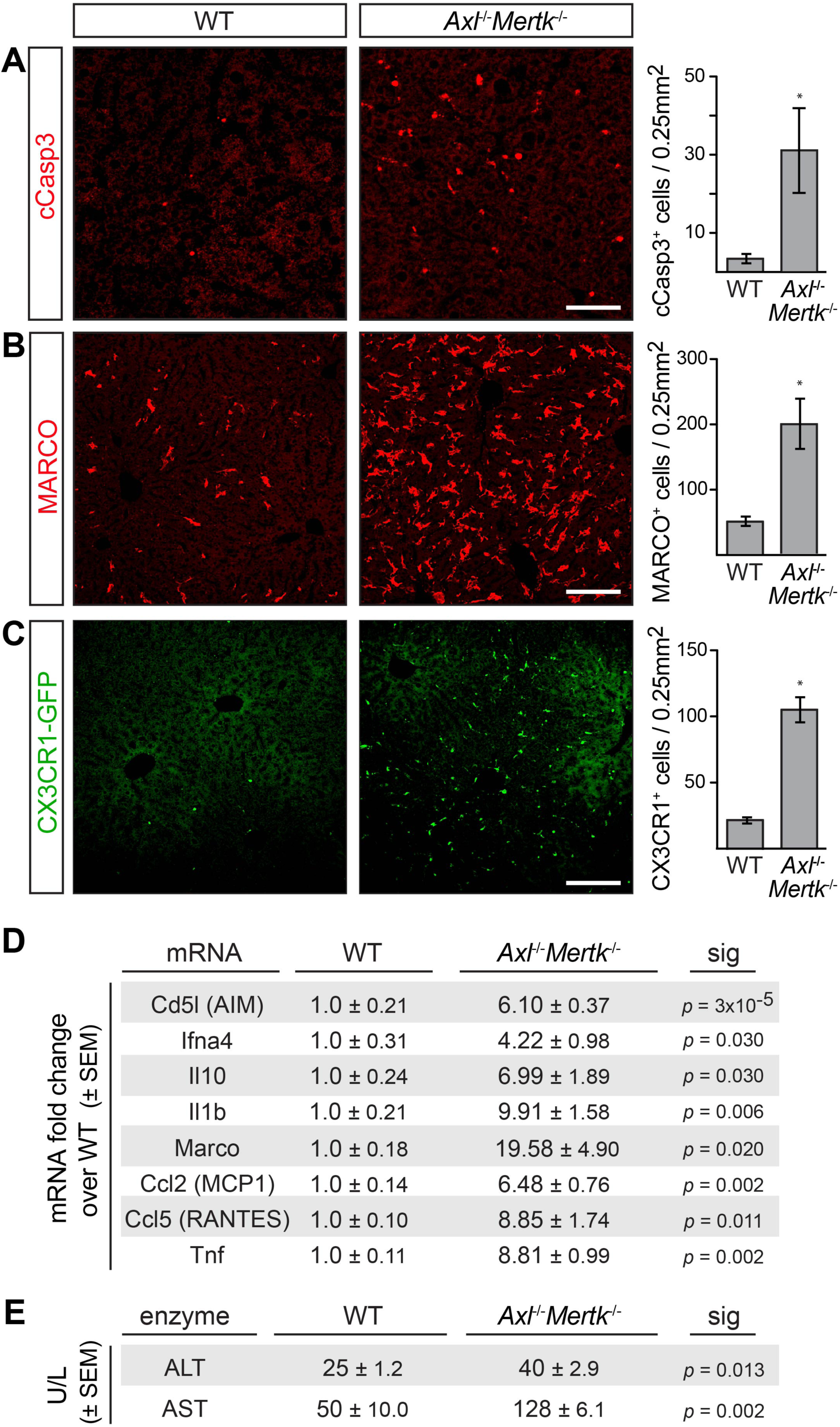
Liver pathology in aged *Axl*^-/-^*Mertk*^-/-^ mice. (A) 9-12 month-old *Axl*^-/-^*Mertk*^-/-^ mice accumulate ∼9-fold more cCasp3^+^ ACs than WT mice. (B) 9-12 month-old *Axl*^-/-^*Mertk*^-/-^ exhibit a ∼4-fold increase in MARCO staining in liver compared to WT. (C) 9-12 month-old *Axl*^-/-^*Mertk*^-/-^*Cx3Cr1*^+/GFP^ mice have ∼5-fold more infiltrating GFP^+^ immune cells in liver compared to *Cx3Cr1*^+/GFP^. (D) Levels of the indicated mRNAs, isolated from 8 month livers, and quantified by qPCR relative to WT. (E) Serum ALT and AST activity from 7 month-old WT and *Axl*^-/-^ *Mertk*^-/-^ mice. (A-C) Representative images from 3-4 mice per genotype. (D-E) Representative data from 2 independent experiments (n = 3-4 mice per genotype). Bars (A-C) 100 µm; * p<0.05.

### Role of TAM receptors in Jo2 and LPS/D-Gal acute injury models

We next asked how TAM receptor mutants would fare in two acute liver injury models – a fulminant hepatitis model based on injection of the Jo2 anti-Fas antibody (42), and an endotoxic shock model precipitated by injection of lipopolysaccharide (LPS) and D-galactosamine (43). A nonlethal intraperitoneal (IP) dose of Jo2 (0.3 mg/kg) did not lead to any change in the appearance of either the WT or *Axl*^*-/-*^ liver at 24 h after treatment, but produced widespread hemorrhage and severe congestion of the sinusoidal space in both the *Mertk*^*-/-*^ and *Axl*^*-/-*^*Mertk*^*-/-*^ liver (Fig. 3A). Similarly, while this treatment yielded few uncleared ACs in WT and *Axl*^*-/-*^ livers, a 15-fold increase in ACs was seen in *Mertk*^*-/-*^ and *Axl*^*-/-*^*Mertk*^*-/-*^ livers (Fig. 3B,C). Serum ALT and AST levels were also elevated specifically in *Mertk*^*-/-*^ and *Axl*^*-/-*^*Mertk*^*-/-*^ mice (Fig. 3D,E). Consistent with these results, the non-lethal Jo2 dose in WT mice led to enhanced hepatic activation of Mer, but not Axl, as assessed by tyrosine autophosphorylation (Fig. 3F). When we used a nonlethal dose of LPS/D-gal, we similarly observed very few ACs in WT or *Axl*^*-/-*^ mice at 16 h after injection, but detected many in both *Mertk*^*-/-*^ and *Axl*^*-/-*^*Mertk*^*-/-*^ mice (Fig. 4A-C). ALT and AST levels were again markedly elevated in *Mertk*^*-/-*^ and *Axl*^*-/-*^*Mertk*^*-/-*^ mice, but not in *Axl*^*-/-*^ mice (Fig. 4D,E). Recovery and regeneration from LPS/D-gal damage was delayed in *Mertk*^*-/-*^ and *Axl*^*-/-*^*Mertk*^*-/-*^ mice, since loci of post-apoptotic necrotic cells and Ki67^+^ proliferative cells were still present in these mice at 7 d after injection (Fig. 4F). Together, these results demonstrated that TAM receptors are required for liver homeostasis, and that Mer specifically is essential for the clearance of apoptotic cells that are produced during acute injury.

**FIG. 3.**
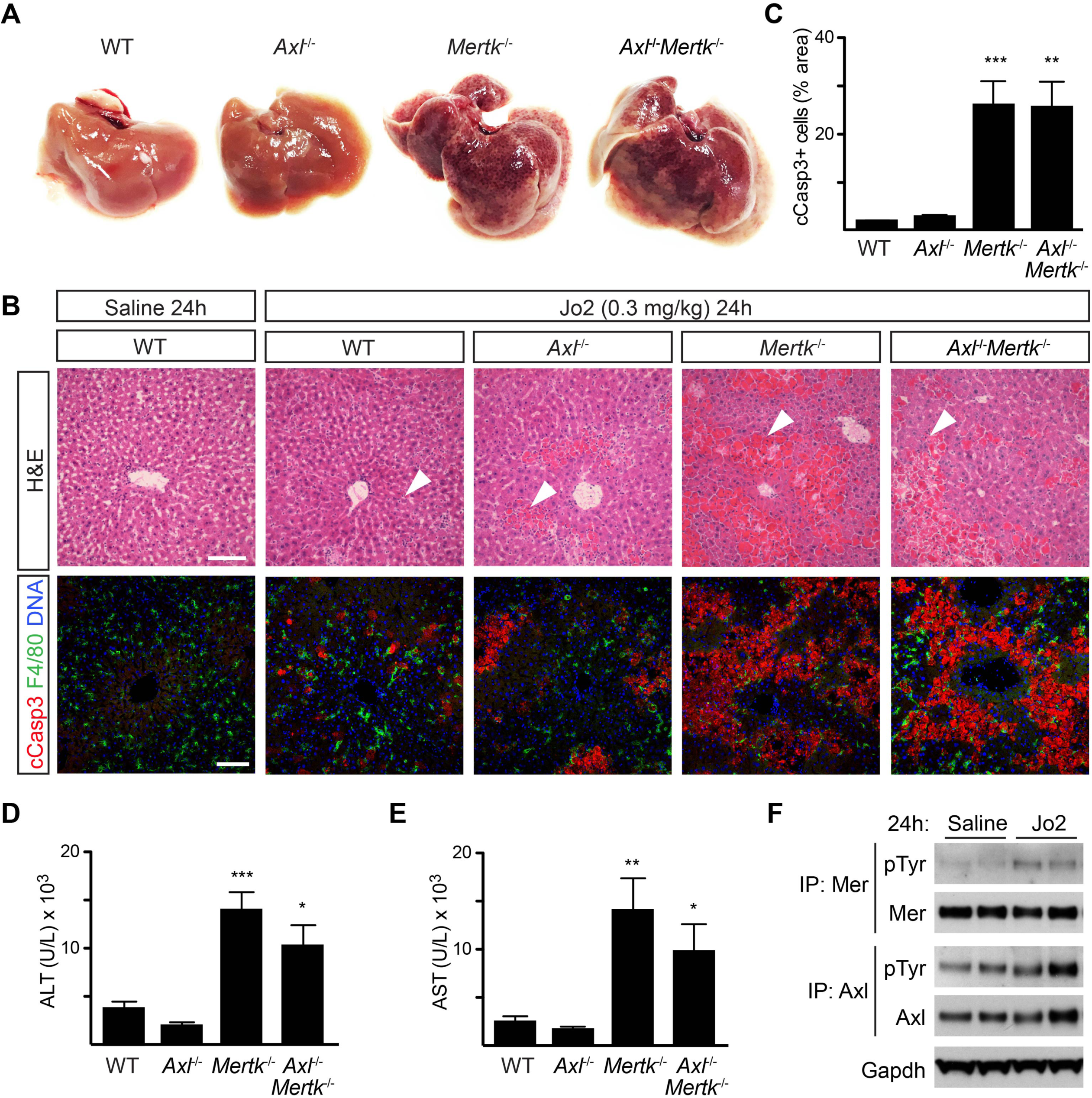
Protective role of Mer in non-lethal Jo2 liver damage model. (A) Extensive hemolysis (mottled darkening) of liver lobes 24 h after a non-lethal (0.3 mg/kg) Jo2 injection in perfused *Mertk*^*-/-*^ and *Axl*^*-/-*^*Mertk*^*-/-*^, but not in *WT* or *Axl*^*-/-*^ mice. (B) Liver sections were analyzed by H&E staining (top row) and immunohistochemistry with indicated antibodies (bottom row). Markedly increased accumulation of apoptotic and necrotic cells is observed in *Mertk*^*-/-*^ and *Axl*^*-/-*^ *Mertk*^*-/-*^ liver. Bars, 100 µm. Arrowheads: damaged tissue, AC accumulation. A,B: Representative images from 3 experiments (n = 3-5 mice per genotype). (C) Apoptotic area quantification demonstrates a ∼14-fold increase in cCasp3^+^ cells. (D,E) ALT/AST serum activity assays. n=7-9 mice per genotype. * p<0.05, **\*\*** p<0.01, *** p<0.001. (F) Liver lysates from WT mice injected with saline or Jo2 were immunoprecipitated for Mer and Axl and immunoblotted with indicated antibodies. Representative image from 2 experiments (n = 2 mice/treatment).

**FIG. 4.**
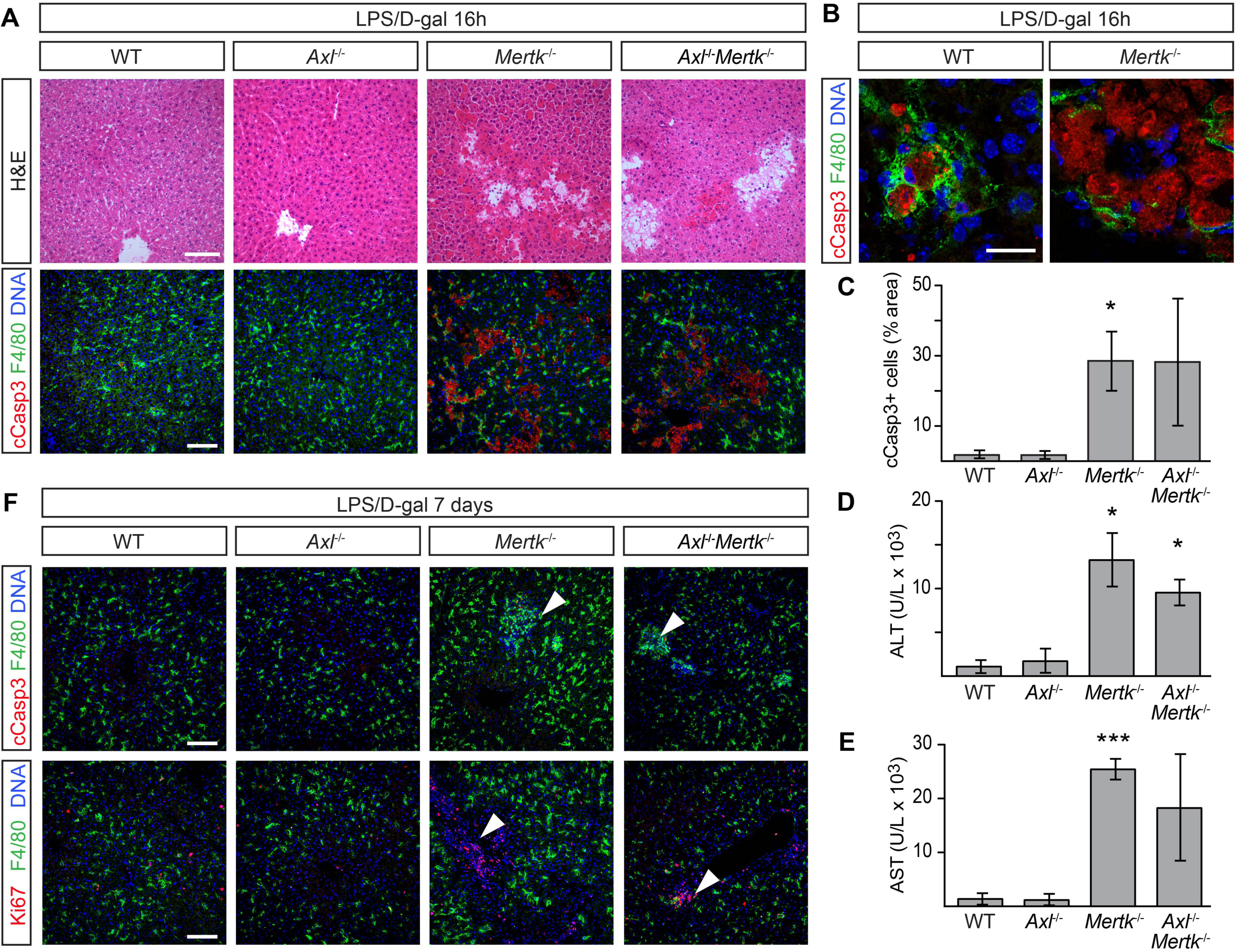
Protective role of Mer in non-lethal LPS/D-gal liver damage model. (A, B) Liver sections were analyzed 16 h after IP injection with 350 mg/kg D-gal and 1 µg/kg LPS by H&E staining and immunohistochemistry with the indicated antibodies. Increased accumulation of apoptotic and necrotic cells can be observed in *Mertk*^*-/-*^ and *Axl*^*-/-*^*Mertk*^*-/-*^. Bars, 100 μm (A), 20 μm (B). (B) shows that ACs are engulfed by F4/80^+^ KCs. (C) Apoptotic area quantification. (D, E) ALT and AST activity assays in sera 24 h post-injection. (F) Liver sections 7 days post injection analyzed by H&E staining and immunohistochemistry with the indicated antibodies. Arrowheads in upper and lower rows represent unresolved loci of condensed, necrotic cCasp3^+^ cells and proliferating Ki67^+^ cells, respectively, specifically in *Mertk*^*-/-*^ and *Axl*^*-/-*^*Mertk*^*-/-*^ livers. Bars, 100 μm. n=3 per genotype, except *Axl*^*-/-*^*Mertk*^*-/-*^ n=2. * p ≤ 0.05, *** p ≤ 0.005.

### TAM receptor expression in KCs is critical for liver physiology

We asked whether these phenomena reflected TAM expression in KCs versus endothelial cells (ECs). KCs are transiently CX3CR1^+^ during their development, but are CX3CR1^-^ in the mature liver (34) (Fig. 1E,F, Fig. 2C), whereas ECs are never CX3CR1^+^. We used a constitutively active *Cx3cr1*^*Cre*^ line, which drives Cre expression early in KC development (34), and crossed this line with conditional floxed *Mertk*^*f/f*^ (9) and *Axl*^*f/f*^ (33) alleles. We also crossed the tamoxifen (Tx)- inducible *Cx3cr1*^*CreER/+*^ line (44), which we have used previously (9), to these same conditional alleles. The *Cx3cr1*^*Cre/+*^*Axl*^*f/f*^*Mertk*^*f/f*^ mice displayed a dramatic reduction in Axl and Mer expression in KCs and peritoneal macrophages, whereas KC expression in Tx-injected *Cx3cr1*^*CreER/+*^*Axl*^*f/f*^*Mertk*^*f/f*^ mice was unaffected (Fig. 5A-C). The *Cx3cr1*^*CreER/+*^ line only drives Cre expression in CX3CR1^+^ cells upon Tx activation, and as noted above, mature KCs are CX3CR1^-^. At 7 months of age, the *Cx3cr1*^*Cre/+*^*Axl*^*f/f*^*Mertk*^*f/f*^ livers displayed elevated levels of many proinflammatory and immunoregulatory mRNAs relative to *Axl*^*f/f*^*Mertk*^*f/f*^ (Fig. 5D). In addition, a non-lethal Jo2 dose led to significantly greater AC accumulation in *Cx3cr1*^*Cre/+*^*Axl*^*f/f*^*Mertk*^*f/f*^ livers than in Tx-treated *Cx3cr1*^*CreER/+*^*Axl*^*f/f*^*Mertk*^*f/f*^ livers (Fig. 5E). These results argue that TAM expression in KCs is critical for the phenotypes that develop during both aging and the response to acute injury.

**FIG. 5.**
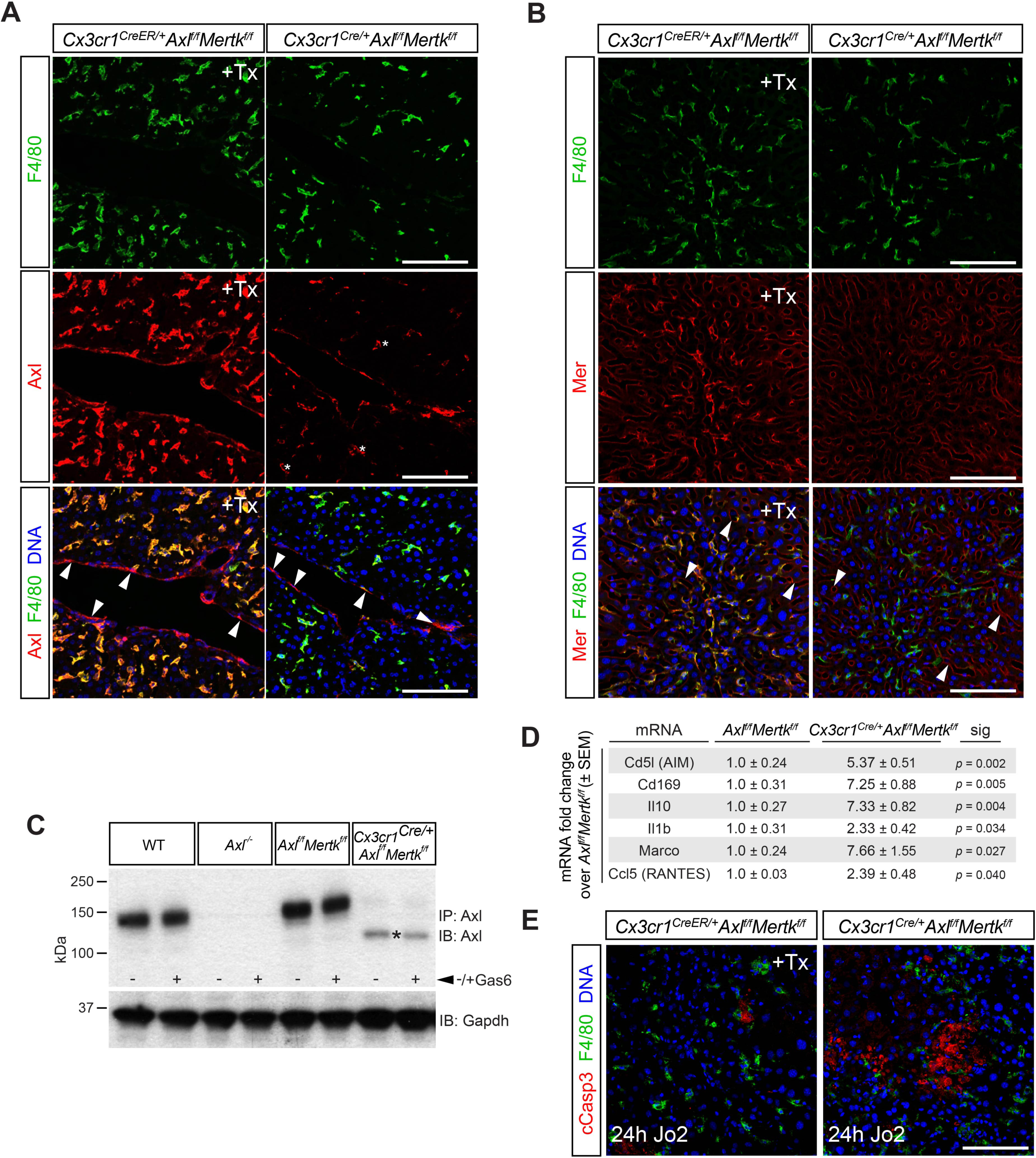
Axl deletion from KCs. (A) Liver sections from tamoxifen-treated (+Tx) *Cx3cr1*^*CreER/+*^*Axl*^*f/f*^*Mertk*^*f/f*^ mice and constitutive *Cx3cr1*^*Cre/+*^*Axl*^*f/f*^*Mertk*^*f/f*^ mice, stained with antibodies to F4/80 (top row, green) and Axl (middle row, red), illustrating *Cx3cr1*^*Cre/+*^-driven loss of Axl from KCs but not ECs in constitutive *Cx3cr1*^*Cre/+*^*Axl*^*f/f*^*Mertk*^*f/f*^ mice, but normal strong Axl expression in KCs in Tx-treated *Cx3cr1*^*CreER/+*^*Axl*^*f/f*^*Mertk*^*f/f*^ mice. Bottom row is merged image of top and middle rows, with DNA visualized by Hoechst 33258 (blue). Note that Axl immunostaining in KCs of the constitutive *Cx3cr1*^*Cre/+*^*Axl*^*f/f*^*Mertk*^*f/f*^ liver is dramatically reduced, but not completely eliminated. Examples of KCs with low residual Axl staining are indicated by asterisks. Axl immunostaining in ECs (arrowheads) persists in both genotypes. (B) Same analyses as in A, except that middle row sections are stained with a Mer antibody. Mer immunostaining in ECs (arrowheads) persists in both genotypes. (C) Peritoneal macrophages from *Cx3cr1*^*Cre/+*^*Axl*^*f/f*^*Mertk*^*f/f*^ mice, +/- treatment with 10 nM Gas6, were immunoprecipitated and blotted for Axl. The low residual level of Axl immunostaining in KCs of these mice (A) may be due to the presence of a low level of a truncated Axl (asterisk) produced by Cre-mediated recombination. (D) Levels of the indicated mRNAs, isolated from 6-7 month livers of the indicated genotypes, and quantified by qPCR. (E) Cleaved Casp3^+^ AC accumulation in the liver 24 h after a non-lethal Jo2 injection in Tx-injected *Cx3cr1*^*CreER/+*^*Axl*^*f/f*^*Mertk*^*f/f*^ mice, in which Axl and Mer expression in KCs is maintained (left), versus *Cx3cr1*^*Cre/+*^*Axl*^*f/f*^*Mertk*^*f/f*^ mice, in which Axl and Mer expression in KCs is lost (right). Bars (A, B, E), 100 μm.

### Role of TAM receptors in acetaminophen-induced acute liver injury

Acetaminophen (APAP)-induced hepatotoxicity is the most frequent cause of acute liver failure worldwide (45, 46). We therefore sought to examine the relative performance of WT, *Axl*^*-/-*^, and *Mertk*^*-/-*^ mouse mutants in a standard model of acute APAP intoxication, which involves overnight (16 hour) fasting and subsequent IP injection of the drug at 300 mg/kg (see Materials and Methods). Although analyses in this model are generally focused on the resolution of inflammation, which occurs between 48 and 72 hours after acetaminophen overdose, we found that studies of APAP-treated mice beyond 48 hours after drug administration were precluded by the remarkably strong phenotype that we observed specifically in *Axl*^*-/-*^ mice. In two independent experimental series, both WT and *Mertk*^*-/-*^ mice were motile and superficially normal at 12, 24, and 48 hours after APAP, but most *Axl*^*-/-*^ mice were very sick and non-motile across all of this period. Examination of the livers of APAP-treated mice at 48 hours after drug administration revealed massive hemorrhage specifically in the *Axl*^*-/-*^ mice (Fig. 6A, B). A typical *Axl*^*-/-*^ APAP-treated liver in situ, seen in 60% of treated mice, is shown in Fig. 6A. Histological staining of liver sections 48 hours post-treatment revealed substantial hemorrhage within the *Axl*^*-/-*^ but not WT or *Mertk*^*-/-*^ liver parenchyma (Fig. 6B). Serum levels of ALT were also markedly elevated at 48 hours after APAP specifically in *Axl*^*-/-*^ relative to WT and *Mertk*^*-/-*^ mice (Fig. 6C), and although APAP intoxication is primarily associated with necrosis, APAP-treated *Axl*^*-/*-^ livers also displayed elevated numbers of cCasp3^+^ cells relative to wild-type (Fig. 6D). Enzymatic activation of the Axl tyrosine kinase always triggers downstream metalloprotease cleavage of the Axl extracellular domain from the cell surface, and the generation of soluble Axl (sAxl) (7, 13, 47). Correspondingly, we measured elevation of circulating sAxl in serum 48 hours after APAP treatment of WT mice (Fig. 6E). We did not detect significant elevation of *Cd5l, Il10, Il1b, Ccl2, Ccl5*, or *Tnf* mRNAs in *Axl*^*-/-*^ versus either WT or *Mertk*^*-/-*^ livers, indicating that the Axl-specific liver damage induced by APAP was not due to specific elevation of these cytokine/chemokine mRNAs (Fig. 6F). Our results with *Axl*^*-/-*^ mice stand in marked contrast to earlier studies of APAP intoxication in *Mertk*^*-/-*^ mice, which resulted in relatively modest changes in necrosis and neutrophil infiltration of the liver only at early (8-24 hour) post-treatment timepoints (48).

**FIG. 6.**
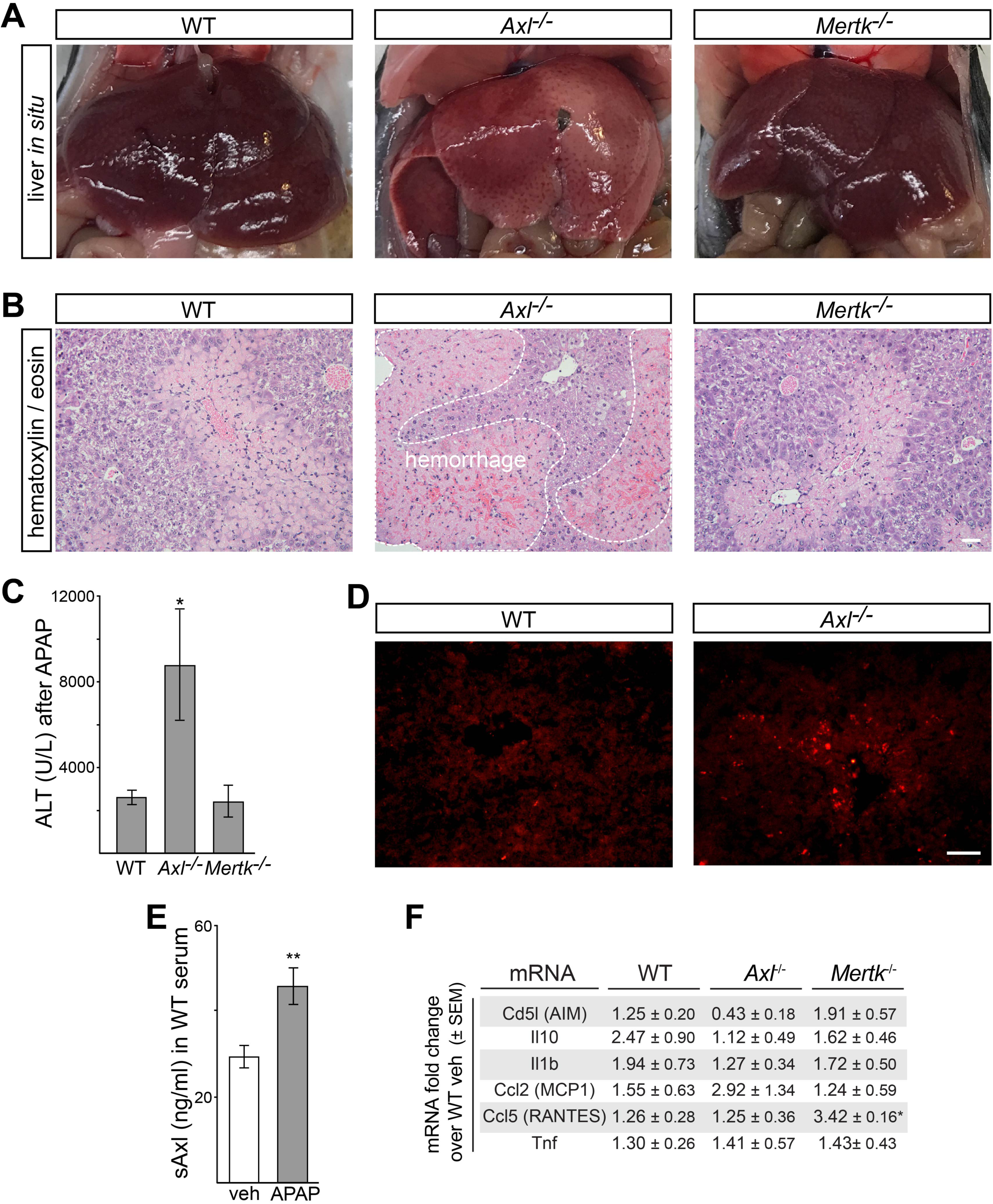
Protective role of Axl in APAP intoxication. (A) Extensive hemorrhage and congestion of liver lobes 48 h after administration of acetaminophen (APAP; 300 mg/kg) to *Axl*^*-/-*^ but not *WT* or *Mertk*^*-/-*^ mice. These *in situ* liver images are from *non-perfused* mice. (B) Liver sections from 48 h APAP-treated mice were analyzed by H&E staining. Markedly increased congestion and blood hemorrhage is observed in *Axl*^*-/-*^ but not *WT* or *Mertk*^*-/-*^ liver. Bars, 100 µm. (C) Measurement of circulating ALT (U/L, units per liter) 48 h after APAP administration in mice of the indicated genotypes. (D) Immunostaining for cleaved Casp3^+^ cells in liver sections of WT and *Axl*^*-/-*^ mice at 48 h after APAP administration. (E) Induction of soluble Axl (sAxl) in WT mice 48 h following APAP. As for all APAP treatments, mice were fasted for 16 h prior to drug administration. (E) Levels of the indicated mRNAs, isolated livers of the indicated genotypes 48 h after APAP treatment, and quantified by qPCR. Bars (B, D) 50 μm.

### Role of TAM receptors in response to lethal liver injury

We also observed TAM regulation of the response to a lethal liver injury – a high (1 mg/kg) dose of Jo2 - but with differences in the relative requirement for Axl and Mer. This dose led to AC accumulation at 2 and 4 hours both in WT and in *Axl*^*-/-*^*Mertk*^*-/-*^ mice, with tissue damage at 2 h was much worse in the latter (Fig. 7A). It resulted in the rapid activation of both Mer and Axl in the WT liver (Fig. 7B), although Axl activation was obscured by metalloprotease cleavage of the Axl ectodomain and consequent reduction of steady-state Axl protein after Axl activation (13) (Fig. 7B). This cleavage resulted in the appearance of elevated soluble Axl extracellular domain (sAxl) in the blood at 2 h after Jo2 injection (Fig. 7C), suggesting that circulating sAxl could serve as a biomarker of acute liver damage. Jo2 lethality was enhanced in *Tyro3*^*-/-*^*Axl*^*-/-*^*Mertk*^*-/-*^ mice relative to WT (Fig. 7D), but also in *Mertk*^*-/-*^ and especially *Axl*^*-/-*^ single mutants (Fig. 7E). This may be related to TAM suppression of stimulus-induced inflammatory cytokine production in dendritic cells and macrophages (12, 49, 50), since we observed higher levels of TNFα (Tnf), type I interferon (Ifn), and IL-1β (Il1b) mRNAs in *Axl*^*-/-*^*Mertk*^*-/-*^ liver relative to WT after Jo2 administration (Fig. 7F). Notably, Ifna4 and Ifnb mRNAs were up-regulated specifically in the *Axl*^*-/-*^ and *Axl*^*-/-*^*Mertk*^*-/-*^ but not the *Mertk*^*-/-*^ liver (Supporting Fig. S1A-J).

**FIG. 7.**
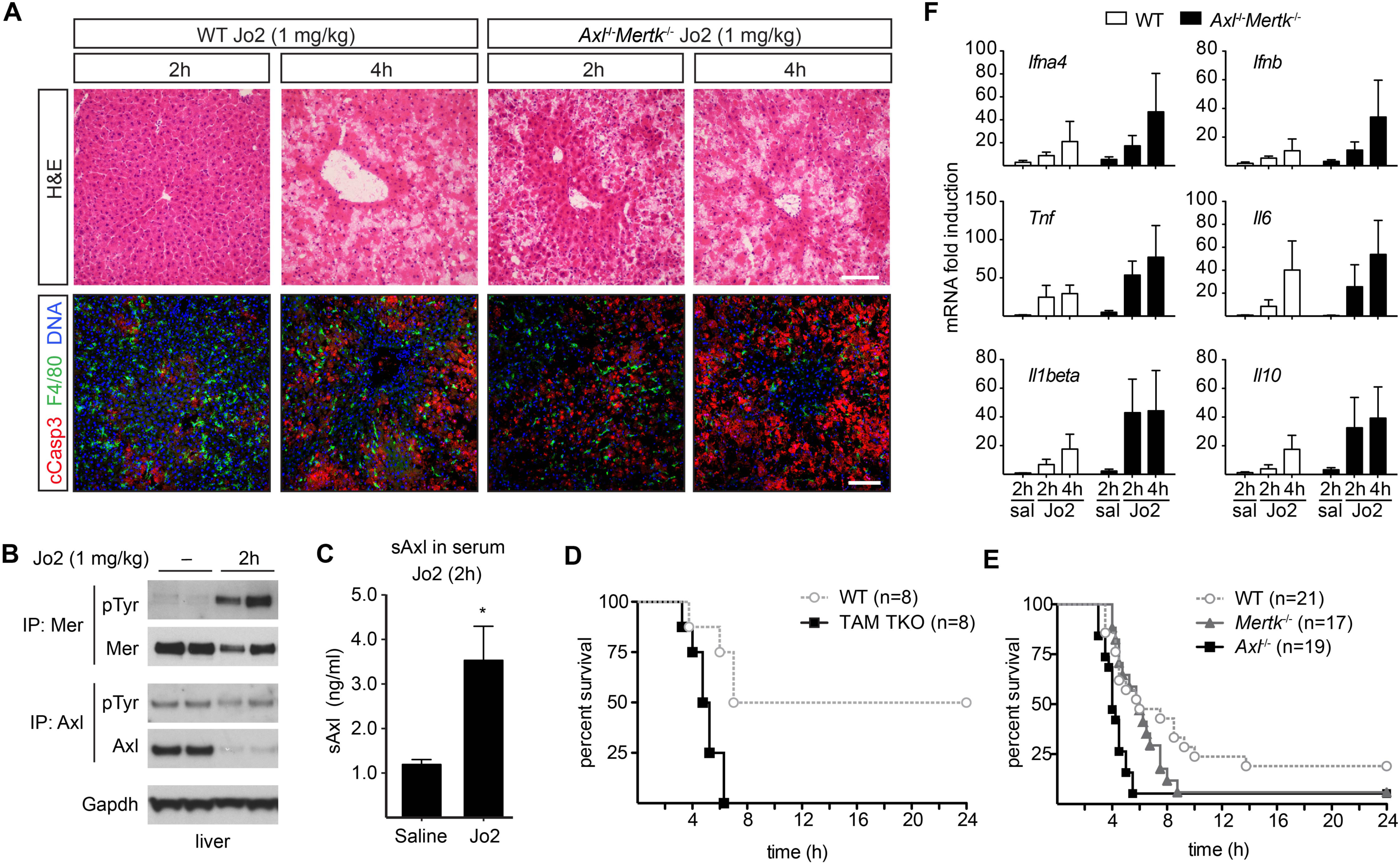
Protective role of Axl and Mer in lethal Jo2 liver damage model. (A) Liver sections at 2 and 4 h after a lethal IP injection (1 mg/kg) of Jo2 were analyzed by H&E staining and immunohistochemistry with indicated antibodies. Increased accumulation of apoptotic and necrotic cells can be observed in *Axl*^-/-^*Mertk*^-/-^. Bars, 100 µm. Representative images from 3 experiments (n = 2-5 mice per genotype.) (B) Liver lysates from WT mice injected with saline or Jo2 (1 mg/kg) were immunoprecipitated for Mer and Axl and immunoblotted with indicated antibodies; n = 2, each lane an individual mouse. (C) sAxl ELISA in serum of saline and Jo2 (1 mg/kg) injected mice; n = 2, * p<0.05. (D) Survival of WT and TAM TKO mice after Jo2 (1 mg/kg) injection. (E) Survival of WT and *Axl*^*-/-*^ and *Mertk*^*-/-*^ mice after Jo2 (1 mg/kg) injection. (F) Expression of indicated inflammatory markers was analyzed by qPCR from liver mRNA samples. (D-F), Representative results from 3 experiments.

### Role of TAM receptors in liver fibrosis

Finally, we examined the role of TAM signaling during carbon tetrachloride (CCl_4_) toxicity - a model of chronic liver damage (51) in which WT, *Axl*^*-/-*^, *Mertk*^*-/-*^, and *Axl*^*-/-*^*Mertk*^*-/-*^ mutants were injected with CCl_4_ three times per week for six weeks. We observed strikingly different results from those seen in the acute injury models: CCl_4_-driven hepatic fibrosis was specifically *enhanced* by Axl signaling (Fig. 8). Collagen deposition was comparable in WT, *Mertk*^*-/-*^, and *Axl*^*-/-*^*Mertk*^*-/-*^ livers, but was *reduced* in the *Axl*^*-/-*^ liver (Fig. 8A, D). Correspondingly, deposition of collagen-associated laminin was also reduced only in the *Axl*^*-/-*^ liver (Fig. 8B, E). In contrast, the phagocytic clearance of ACs was, as seen for the acute injury models, again entirely dependent on Mer (Fig. 8C, F), which may account for the lack of fibrosis protection seen in the *Axl*^*-/-*^*Mertk*^*-/-*^ liver. Long-term CCl_4_ exposure led to a substantial up-regulation of hepatic Axl and Gas6 (Fig. 8G, H). Axl up-regulation was modestly associated with KCs and activated stellate cells (Supporting Fig. S2A-C), but most prominently with loci of infiltrating monocytes (Supporting Fig. S2D), which play critical roles in the progression of both acute and chronic liver disease (52). Our conclusions from the CCl_4_ fibrosis models are in substantial agreement with previously published analyses (24).

**FIG. 8.**
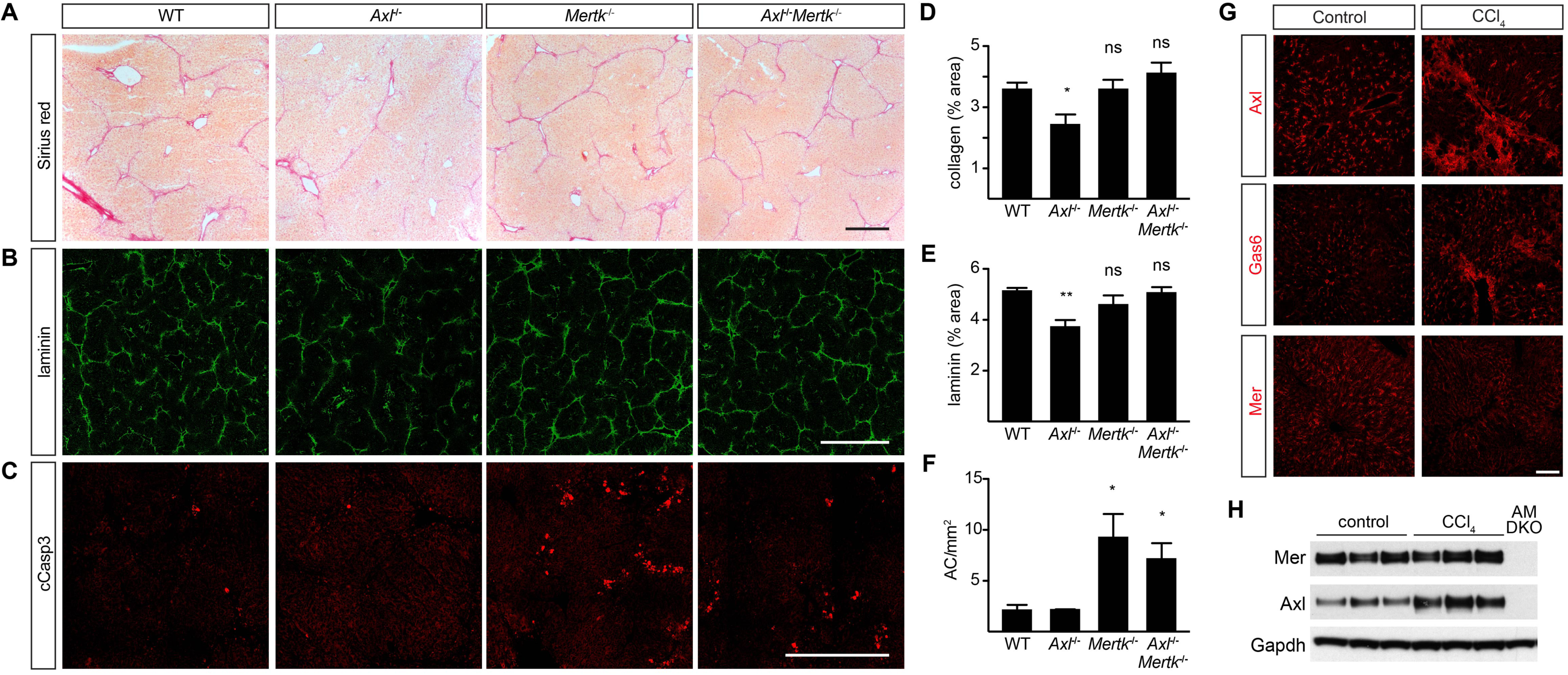
Axl promotion of CCl_4_–induced liver fibrosis. (A) Sirius red staining, showing collagen deposition, of liver sections from mice of the indicated genotypes, injected with CCl_4_ three times per week for 6 weeks. (B) Laminin staining of similar sections. (C) Cleaved caspase 3 (cCasp3) staining of similar sections. (D-F) Quantification of the results in A-C, respectively. (G) Upregulation of Axl and Gas6, but not Mer, in fibrotic liver. (H) Western blot showing increased expression of Axl in fibrotic liver. Bars (A-C) 0.5 mm, (G) 100 µm. Representative images from 6 mice per genotype. * p<0.05, ** p<0.01.

## Discussion

Together, our results demonstrate that TAM RTKs are critical regulators of hepatic physiology and homeostasis. The livers of normally aged *Axl*^*-/-*^*Mertk*^*-/-*^ mice display substantially elevated AC accumulation, pronounced immune activation, and marked hepatic tissue damage relative to their WT counterparts, in the absence of any experimental perturbation. These findings indicate that sustained Axl and Mer signaling throughout adult life is required for healthy aging of the liver.

Significantly, Mer alone is required for the phagocytosis of ACs that are generated in settings of acute liver damage induced by Jo2, and Mer and Axl act in concert to suppress inflammatory responses in the liver, as they do in other organs. These findings, together with our observation that Kupffer cells are a particularly important locus of Mer action, are consistent with the fact that relatively high steady-state expression of Mer is a defining feature of phagocytic tissue macrophages throughout the body (53) and with the demonstration that Mer is absolutely required for the efficient clearance of ACs by these cells (9, 13, 41, 54, 55).

In striking contrast to the Jo2 acute injury model, the most dramatic effects of TAM deletion in the APAP intoxication model are not seen with *Mertk*^*-/-*^ mutants. While these *Mertk* mutants display a ∼1.4-fold increase in necrotic cells and a ∼2-fold increase in activated neutrophils 24 hours after APAP administration (23), many *Axl*^*-/-*^ mice were at or near death from hepatic congestion and hemorrhage by 48 hours after the same APAP treatment. It is important to note that these phenotypes - sinusoidal congestion and hemorrhage - are common features of acetaminophen induced liver injury. Previous studies have revealed that APAP treatment results in excess generation of the fibrinolytic enzyme plasmin, which in turn leads to matrix degradation that detaches sinusoidal endothelial cells from the underlying basement membrane (56). This process, which most likely results from plasmin-dependent activation of matrix metalloproteinases, disrupts sinusoidal blood flow leading to congestion and hemorrhage. Although the mechanism by which Axl might limit plasmin-dependent loss of sinusoidal integrity is unknown, it is interesting to note that this RTK is highly expressed in sinusoidal endothelial cells. It is also possible that Axl activation in Kupffer cells normally stimulates the production of factors that might limit plasmin generation or activity.

The co-expression of Mer and Axl together in Kupffer cells, which is also seen in red pulp macrophages and select other macrophage populations but not in tissue macrophages generally, may account for the concerted immunosuppressive action of these two receptors in the liver. Immunosuppression by KCs is critical, since the liver is continuously exposed to endotoxin and other microbial products present in the portal circulation (57). In the absence of TAM-driven immunosuppression, liver inflammation and tissue damage progress rapidly with age.

Soluble Axl ectodomain (sAxl) has been reported to be an accurate biomarker of cirrhosis and the development of hepatocellular carcinoma (58). Our results provide a mechanistic explanation for sAxl release subsequent to Axl activation in the presence of ACs. In settings of cirrhosis, this activation may occur not only in KCs, but also in infiltrating monocytes, activated stellate cells, and even vascular endothelial cells.

In contrast to the beneficial TAM effects seen in acute injury, Axl signaling is deleterious during chronic hepatic fibrosis, as it promotes scarring. This latter finding is consistent with earlier work demonstrating improvement of CCl_4_-induced steatohepatitis and fibrosis in *Gas6*^*-/-*^ and *Axl*^*-/-*^ mice (24, 59). It has previously been reported that HSCs elevate Axl expression upon activation (24, 60), and studies have indicated that AC phagocytosis by HSCs activates them and stimulates collagen production (61, 62). Our results suggest that Axl expression by KCs, activated stellate cells, and infiltrating monocytes together contribute to the deleterious role of Axl in hepatic fibrosis, although the relative contribution of each cell type remains to be determined.

Many different TAM-targeting agents, including biologics and small molecule kinase inhibitors, are currently in pre-clinical development for the treatment of multiple types of cancers, and several have entered clinical trials. Our results have obvious and immediate implications for the potential re-purposing of these agents to the treatment of liver disease, as they demonstrate divergent activities for Mer and Axl in distinct settings of acute and chronic liver injury. The most salient of these relates to specificity: Axl inhibitors used to treat fibrosis must not perturb the Mer activity required for the response to acute liver injury; and correspondingly, Mer agonists used to treat acute injury must not stimulate the Axl activity that promotes fibrosis.

## Acknowledgements

This work was supported by grants from the NIH (R01 NS085296, R01 AI101400, and P30CA014195), the Leona M. and Harry B. Helmsley Charitable Trust (#2012-PG-MED002), the Nomis, H. N. and Frances C. Berger, Fritz B. Burns, and HKT Foundations (to G.L.); and by postdoctoral fellowships from the Marie Curie International Outgoing Fellowship Program (to P.G.T), the Nomis Foundation (to A.Z.), and the Fundación Alfonso Martín Escudero (to L.J.G.). We thank Joe Hash for excellent technical assistance.

## Supporting Material for

**Fig. S1.**
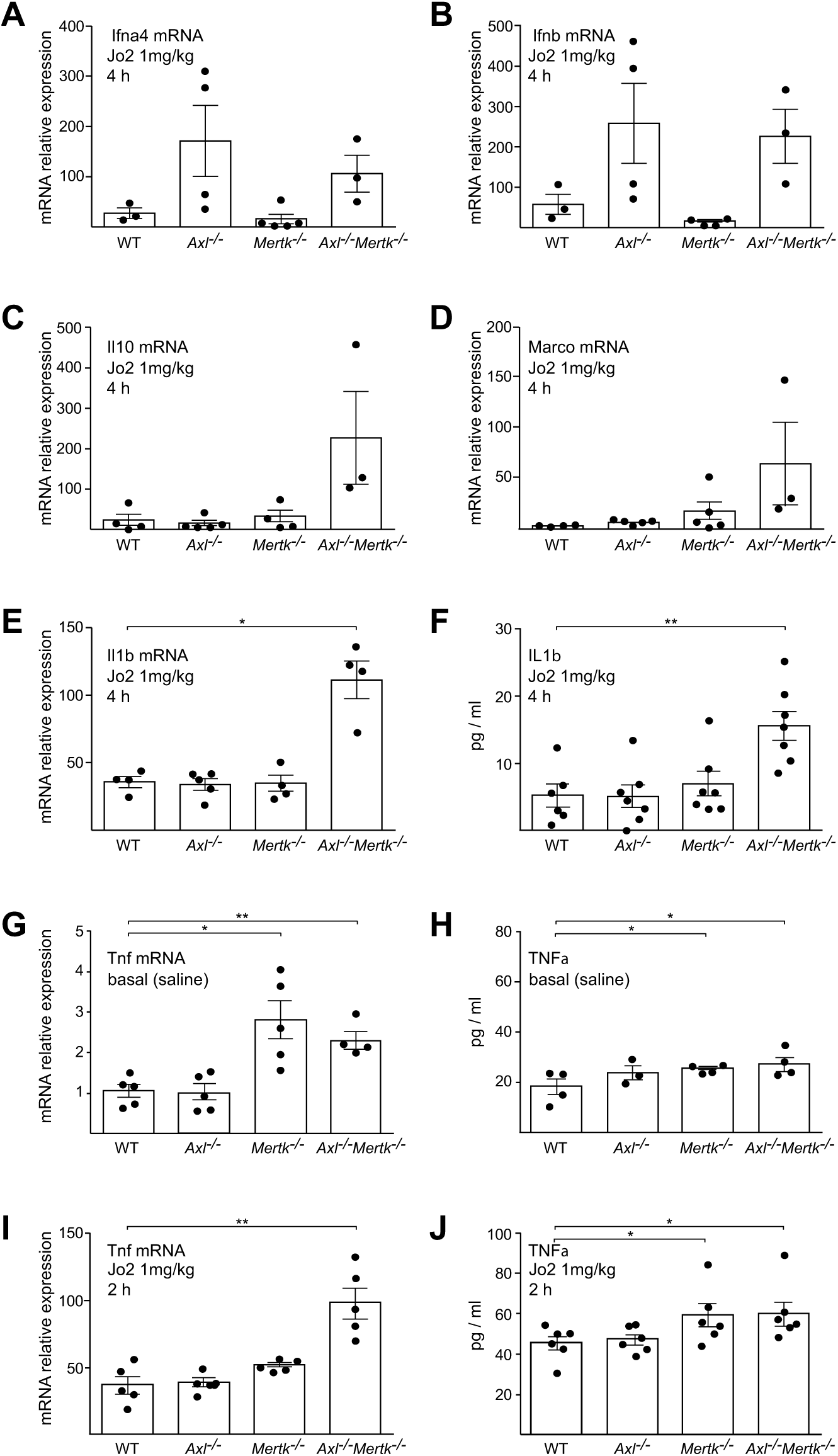
Hepatic cytokine responses to high (lethal) dose Jo2. The indicated mRNAs (A-E,G,I) and proteins (F,H,J) were measured by qPCR and ELISA, at 2 or 4 hours (as indicated) after an IP injection (1 mg/kg) of Jo2 into mice of the indicated genotypes. mRNAs for qPCR were isolated from liver; proteins for ELISA were from serum. Data points are results from separate mice; error bars are ± S.E.M. * p ≤ 0.05, ** p ≤ 0.005.

**Fig S2.**
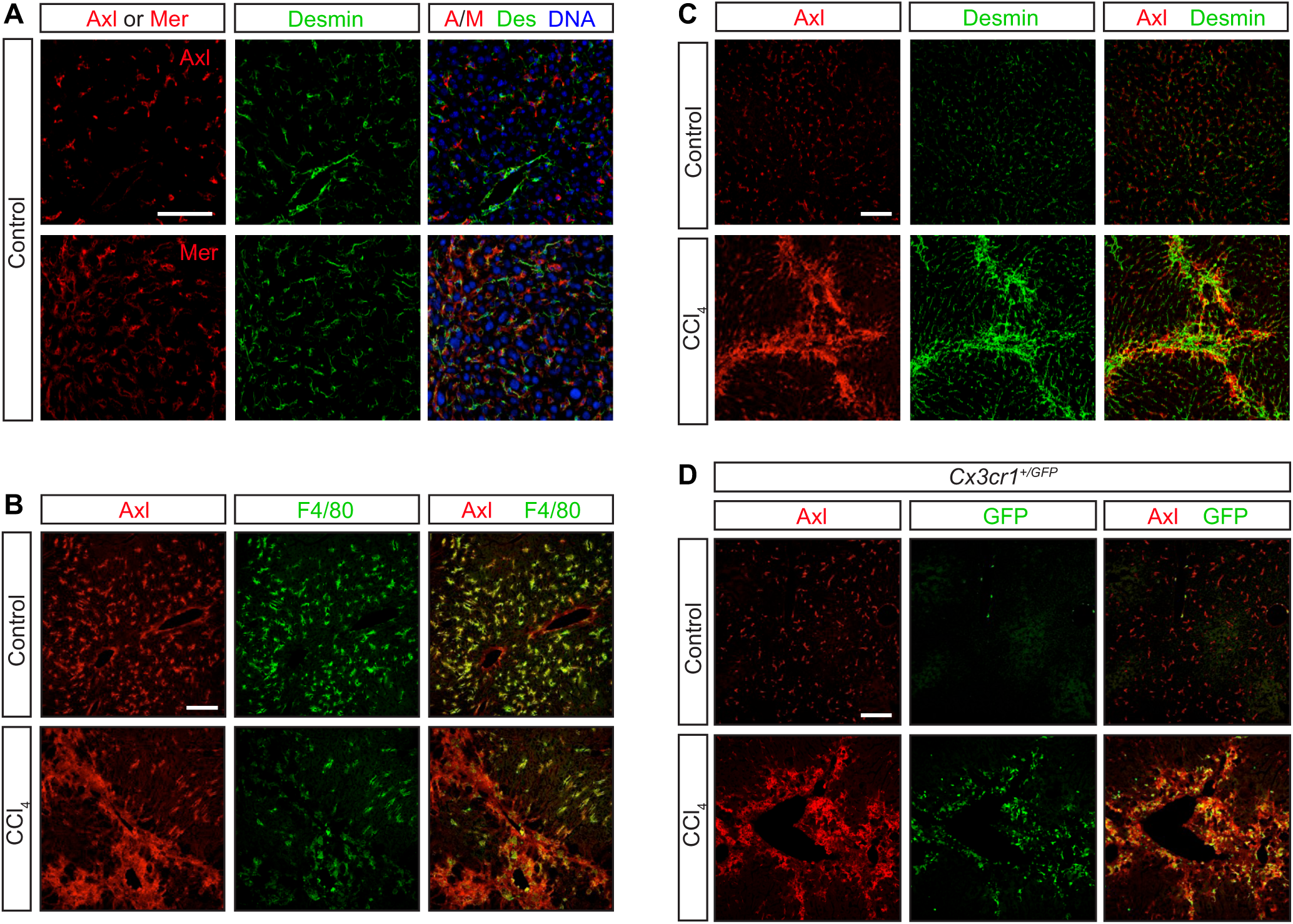
Axl expression in CCl_4_ liver fibrosis model. WT and *Cx3cr1*^*+/GFP*^ mice were injected with CCl_4_ three times per week for 6 weeks. Neither Axl nor Mer co-localizes with the hepatic stellate cell marker Desmin under basal conditions – prior to CCl_4_ treatment (**A, C**, control). As is also shown in Fig. S1A and B, these receptors are instead prominently expressed in F4/80^+^ KCs (**B**). Following CCl_4_ treatment, however, partial overlap between Desmin^+^ and Axl^+^ cells is observed (**C**), although a stronger overlap is seen between Axl and infiltrating CX3CR1^+^ cells (**D**). Bars, 100 µm.

**Supporting Table S1.**
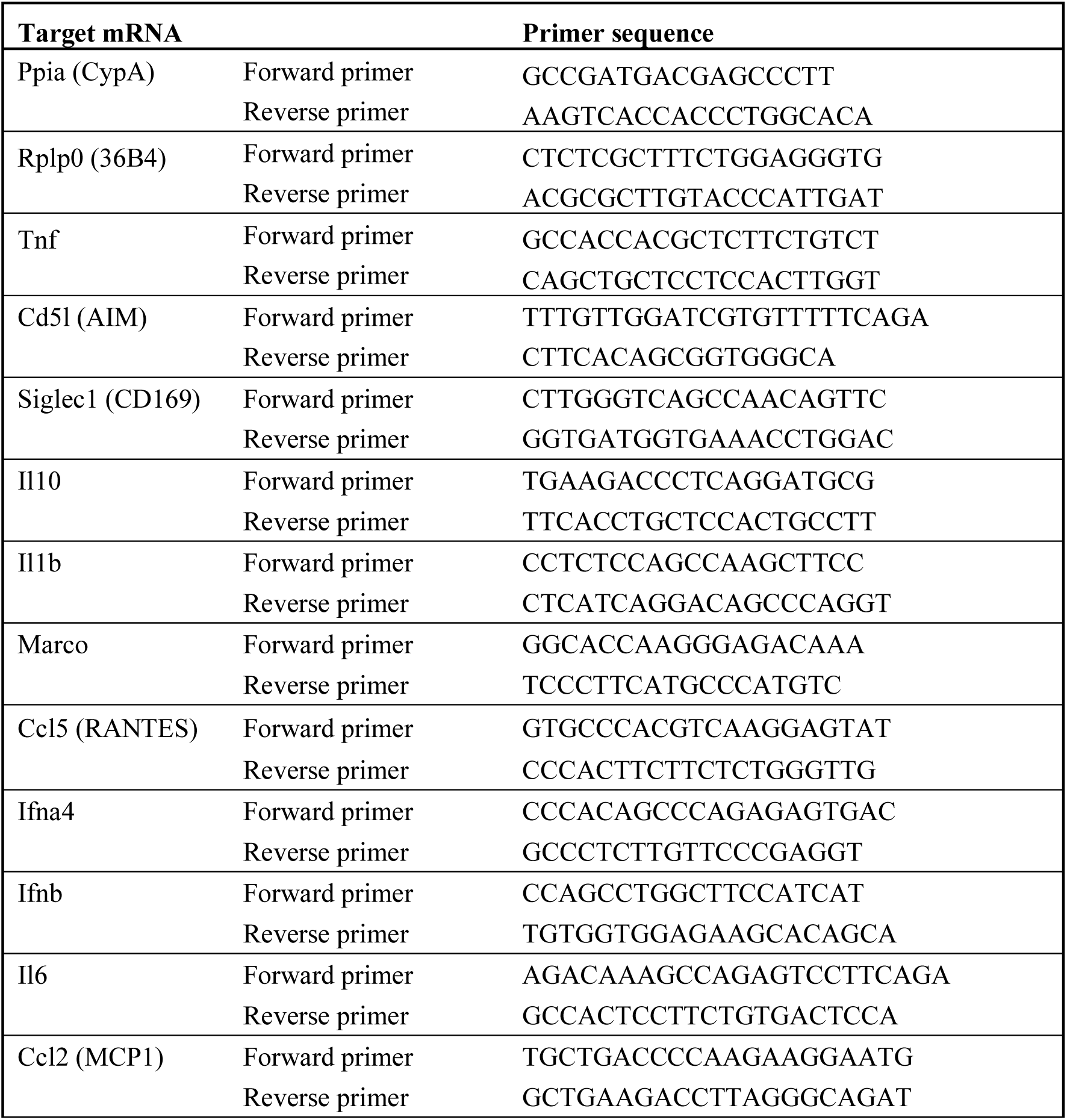

